# Temporal attention and oculomotor effects dissociate distinct types of temporal expectation

**DOI:** 10.1101/2025.03.04.641562

**Authors:** Aysun Duyar, Marisa Carrasco

## Abstract

Temporal expectation–the ability to predict when events occur–relies on probabilistic information within the environment. Two types of temporal expectation, *temporal precision*, based on the variability of an event’s onset, and *hazard rate*, based on the increasing probability of an event with onset delay, interact with temporal attention–ability to prioritize specific moments– at the performance level: Attentional benefits increase with precision but diminish with hazard rate. Both temporal expectation and temporal attention improve fixational stability; however, the distinct oculomotor effects of temporal precision and hazard rate, as well as their interactions with temporal attention, remain unknown. Investigating microsaccade dynamics, we found that hazard-based expectations were reflected in the oculomotor responses, whereas precision-based expectations emerged only when temporal attention was deployed. We also found perception-eye movement dissociations for both types of temporal expectation, yet attentional benefits in performance coincided with microsaccade rate modulations. These findings reveal an interplay among distinct types of temporal expectation and temporal attention in enhancing and recalibrating fixational stability.

## Introduction

The world is dynamic and stochastic, yet it possesses predictable elements that enable us to develop expectations. Temporal expectation, the ability to predict when events occur improves perception in vision (Rohenkohl et al., 2012), audition (Bueti & Macaluso, 2010), and touch (Badde et al., 2020), facilitates actions (Thomaschke & Dreisbach, 2013), alters timing of oculomotor responses (Akdogan & Balci, 2016; Tal-Perry & Yuval-Greenberg, 2021), and modulates neuronal activity at various levels within sensory processing and action pathways (review, Nobre & van Ede, 2023).

Our brain utilizes temporal structures within the environment to develop temporal expectations (Nobre et al., 2007, 2018). The passage of time characterizes <u>hazard rate</u>–the increasing probability of an event, conditional on it not yet having occurred (Coull, 2009; Grabenhorst et al., 2021; Duyar, Denison & Carrasco, 2023)–and the variability of an event’s onset within a time window characterizes <u>temporal precision</u> (Rolke & Hofmann, 2007). These two types of temporal structures yield expectations with different temporal scales: hazard rate is robust to immediate changes whereas temporal precision is computed over longer timescales (e.g., within experimental trials or blocks, respectively).

In the auditory system, hazard rate and temporal precision are partially overlapping and separable at the neural level based on latency, computation, and hierarchical processing stage. The effects of hazard rate on auditory stimulus-evoked MEG responses emerge faster and at lower levels of the hierarchical processing than those of temporal precision. Both effects originate in the early auditory regions, but only temporal precision correlates with activity in the inferior parietal cortices (Todorovic & Auksztulewicz, 2019). Such commonalities and differences are unknown at the oculomotor level.

Temporal attention is another fundamental cognitive process, which facilitates allocation of the brain’s limited resources across time. It helps us prioritize information at specific moments, regardless of predictability (review, Denison, 2024). Temporal attention mediates the effects of temporal expectation at the neural level (Woldorff, Hackley & Hillyard, 1991; Näätänen, 2011; Todorovic et al., 2015). For example, visual temporal attention modulates anticipatory effects in MEG responses (Denison et al., 2024) and interacts with expectation at the behavioral level. In particular, attentional benefits on performance increase with temporal precision, but decrease with hazard rate (Duyar, Ren & Carrasco, 2024). But it is unknown whether temporal attention distinctly interacts with these two types of expectation at the oculomotor level.

Microsaccades (MS), the fastest and largest of fixational eye movements, provide an online metric throughout experimental trials. MS rate follows a typical temporal profile (Martinez-Conde et al., 2009, 2013; Rolfs, 2009). Whereas initially MS were considered as reflexive eye movements, Kowler’s research revealed that they mirror broader aspects of visual perception (reviews: Collewijn & Kowler, 2008; Kowler & Collewijn, 2010; Kowler, 2011), including cognitive factors such as prediction (He & Kowler, 1989) and attention (Kowler, 2024). Typically, there are three stages of the MS rate within an experimental trial (**Fig 1C**): (1) Pre-stimulus endogenous anticipatory reduction, which occurs before an expected brief visual, auditory, or tactile stimulus (Betta and Massimo, 2006; Dankner et al., 2017; Amit et al., 2019; Abeles et al., 2020; Badde et al., 2020). This reduction becomes more pronounced with temporal attention (Denison, Yuval-Greenberg & Carrasco, 2019; Palmieri, Fernandez & Carrasco, 2023), but becomes weaker following a delayed visual event (Tal-Perry & Yuval-Greenberg, 2023; Duyar & Carrasco, 2024). (2) Post-stimulus reflexive exogenous inhibition (Rolfs, Kliegl, Engbert, 2008), which indexes conscious perception (White & Rolfs, 2016) and temporal expectation (Badde et al., 2020). (3) Rebound to baseline, which depends on endogenous inhibition via expectation (Badde et al., 2020), but effects of temporal attention are lacking (Denison et al., 2019; Palmieri et al., 2023). Precise timing of MS is not only modulated by stimulus physical characteristics (Bonneh, Adini, Polat, 2015), but also by task difficulty (Ezzo et al., 2025) and temporal attention (Denison et al., 2019). Temporal attention shifts the timing of both pre-stimulus and post-stimulus microsaccades earlier for temporally predictable targets. However, how temporal attention and different types of expectation work together to recalibrate oculomotor dynamics and stabilize gaze under temporal uncertainty is yet to be investigated.

**Figure 1.**
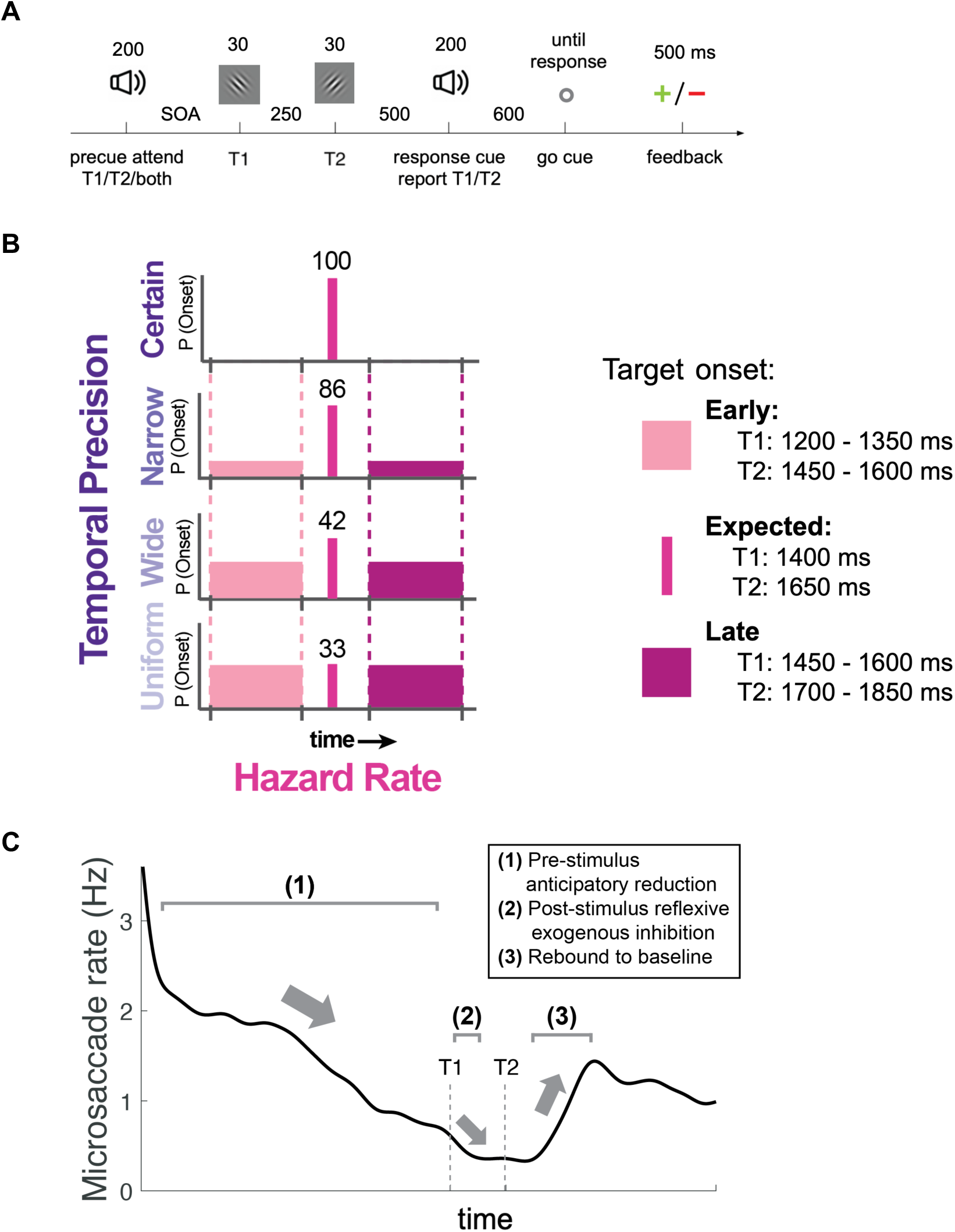
**A.** Experimental protocol (reported in Duyar, Ren & Carrasco, 2024). Attention was manipulated via the precue, and expectation was manipulated via the SOA between the precue and the targets. Observers reported the orientation (clockwise/counterclockwise relative to the reference axis) of the target indicated by the response cue at the end of the trial, with feedback provided based on accuracy. **B.** Hazard Rate was manipulated via within-trial target delay (Early, Expected, Late onsets), and Temporal Precision was manipulated via within-session onset variability (Certain, Narrow, Wide, Uniform). **C.** Typical MS rate profile.

Although perception and eye movements usually co-occur (Vishwanath & Kowler, 2003; Kowler, 2011; Spering & Montagnini, 2011) perceptual reports and eye movements have been dissociated (e.g, Spering, Pomplun & Carrasco, 2010; Spering & Carrasco, 2012; Glasser & Tadin, 2014; Vetter et al. 2019). Recently we found that temporal expectations based on the preceding trial’s stimulus timing is reflected in oculomotor dynamics, but it does not affect the specific time when attention benefits performance (Duyar & Carrasco, 2024). To further our understanding of perception-action dissociations (Spering & Carrasco, 2015; Carrasco & Spering, 2024) here we investigate the eye-tracking dataset of a recent study in which we showed that effects of temporal attention on performance interact in an opposite manner with temporal precision and hazard rate (Duyar et al., 2024). Attentional benefits increase with temporal precision, whereas they decrease with hazard rate. Whether these perceptual effects are also reflected in the microsaccade dynamics is unknown.

Here, we ask: 1) Do hazard rate and temporal precision differentially modulate MS?; 2) Does temporal attention distinctly modulate hazard rate and temporal precision in MS dynamics?; 3) Are perceptual interactions of attention and expectation reflected in MS dynamics, or is there a perception-eye movement dissociation?

## METHODS

### Dataset

We analyzed the eye-tracking data of a previous study in which we investigated behavioral effects on the interaction between temporal attention and expectation (Duyar et al., 2024). The observers, apparatus, stimuli and experimental protocol were identical to that study.

### Observers

Sixteen observers (10 females, 6 males, aged 22-34, M=27, SD=3.445) participated in the study. Two observers were removed from the temporal precision analysis due to corrupted raw eye files for at least one of four conditions; thus, we analyzed the data from the remaining 14 observers (8 females, 6 males, aged 22-34, M=26.786, SD=3.599). For the hazard rate analysis we included all observers because we had eye files for all conditions relevant to that analysis.

All observers had normal or corrected-to-normal vision. The research was performed in accordance with the Helsinki declaration and the experimental protocol was approved by the New York University Institutional Review Board.

### Apparatus

Stimuli were generated using an Apple iMac (3.06 GHz, Intel Core 2 Duo) and MATLAB 2012b (Mathworks, Natick, MA, USA) using Psychophysics Toolbox (Brainard, 1997; Kleiner et al., 2007), presented on a gamma-corrected CRT monitor (1280 x 960 resolution, 100 Hz refresh rate). The viewing distance was 57 cm, and the observers’ head movements were restrained using a chin and headrest. An infrared eye tracker Eyelink 1000 (SR research, Ontario, Canada) was used for recording eye position throughout the experiment. Online eye tracking was implemented to maintain central fixation (1°) throughout the trials. Eye tracking data was recorded at 1000 Hz. Five-point-grid calibration was performed at the beginning of each session, and was repeated within the session when necessary. In case of a fixation break due to blink or ≥1° deviation from the center of the screen, the trial was aborted and repeated at the end of the corresponding experimental block. Observers could blink or make larger saccades during the response window and the intertrial interval.

### Stimuli

Stimuli were displayed on a uniform medium gray background. A central fixation circle subtended 0.15°, and four black circles (0.2°) were displayed at corners of a central imaginary square (2.2° side) as placeholders. Two Gaussian-windowed (SD=0.3°), 100% contrast sinusoidal gratings with a random phase were presented at the center sequentially in each trial. Each grating was tilted clockwise or counterclockwise from the horizontal or vertical axis, independently. The amount of tilt was titrated separately for each target interval for each observer.

The attentional precues were 200 ms auditory tones, presented through speakers. The valid precue was a sinusoidal wave (T1: 800 Hz, T2: 400 Hz), and the neutral precue was a complex waveform that is a combination of a range of frequencies (50-400 Hz). Response cues were identical to the valid precues to instruct the observers to report the first (T1) or the second (T2) target.

### Experimental Procedure

The experimental protocol was reported in detail in Duyar et al. (2024). The experiment was developed by combining a temporal attention (Denison et al., 2017, 2021) and a temporal expectation (Todorovic & Auksztulewicz, 2019) protocol. The experimental protocol is shown in **Fig 1A**.

#### Expectation Manipulations

Two types of temporal expectation –temporal precision and hazard rate– were manipulated (**Fig 1B)**: <u>Temporal precision</u> via within-session target onset variability. The most expected time point (mean point of each temporal distribution) was kept constant (1400 ms for T1, and 1650 ms for T2 relative to the precue) across precision levels, and the probability of stimulus appearance at the expected moment increased with precision: uniform (33%), wide (42%), narrow (86%), certain (100%) conditions. Hence, the variability of target onset was systematically decreased as the precision increased. The <u>hazard rate</u> was manipulated in the low-precision conditions via within-trial target onset delay (early-expected-late). Within the low precision experimental sessions, the targets could appear earlier or later than the expected moment. In a trial with low temporal precision, the conditional probability of the targets’ appearance at each time point increased given that they had not appeared yet, resulting in a lower hazard rate at the earlier than expected moments.

#### Attention Manipulations

Temporal attention was manipulated via auditory precues. The precue instructed observers which target to attend, and the response cue which target’s orientation to report. The precue either instructed observers to attend to the first (T1) or the second (T2) target, or it was uninformative (neutral/baseline) regarding which targets would be behaviorally relevant. The precues corresponding to T1 or T2 were 100% valid, thus the response cue was identical to the precue in those trials. In the neutral trials, the response cue indicated either target with 50% probability.

#### Discrimination Task

The task was to report the orientation (clockwise/counterclockwise) of the target that was indicated by the auditory response cue (T1 or T2). The stimuli orientations were independently selected to have a horizontal or vertical reference axis from which they were slightly offset. Visual feedback (green + or a red -) was presented after each trial based on whether the observers gave a correct or incorrect response. The average accuracy was presented at the end of each block.

#### Training and titration procedures

All observers completed a 60-trial training block in the beginning of the first session to familiarize themselves with the attentional precues and the task. In the beginning of each session/day, the observers completed a titration protocol of 128 trials. We used Best PEST procedure (Liberman & Pentland, 1982), which adjusted the orientation tilt, to attain 75% orientation discrimination accuracy separately for each target in the Neutral condition. Neutral performance was monitored throughout the experiment and, if needed, additional adjustments to the tilt were automatically made after each block. Titration procedure was conducted with constant target timing (**Fig 1B**, temporal precision=certain). Observers were explicitly informed regarding the temporal precision of the session.

### Eye-tracking data preprocessing and microsaccade detection

Each observer completed 3568-4256 trials in total (M=3689, SD=245.773). Due to technical problems the eye data for some blocks was not saved. We recovered ∼90% of the eye-tracking data from 16 people for the hazard rate analysis (M=3291.44, SD=425.36), and from 14 people for the precision analysis (M=3368, SD=393.470).

We used an established velocity-based algorithm to detect MS within each experimental trial (Engbert & Kliegl, 2003). For each trial, we transformed raw eye position data into a 2D velocity space. A trial-specific 2D threshold was then determined as an ellipse centered on the median velocity in both horizontal and vertical directions. An MS was detected when the eye movement velocity exceeded this threshold for at least 6 ms.

### Microsaccade rate timecourse

MS rate timecourses were calculated for each observer, separate for each attentional precue and expectation condition. For each experimental condition, we averaged the number of MS per time samples across all trials, multiplied these values by the sampling rate, and then smoothed across time by applying a 50-ms sliding window.

We performed a series of nonparametric cluster-based permutation tests to compare MS rate timecourses and identify the intervals when they were significantly different (Maris & Oostenveld, 2007). We used two-tailed dependent samples tests (α = 0.05, 2000 permutations) and implemented Bonferroni correction on the identified clusters to account for multiple comparisons. These tests are widely used in MS research, and provide useful information regarding when rate timecourses differ between two conditions within a time period (e.g., Liu, Nobre & van Ede, 2022, Palmieri et al., 2023).

### Microsaccade rate rebound inflection points

As a follow-up to the cluster-based permutation tests, we zoomed into the MS rate window to analyze the timing of MS rate rebound. We estimated the inflection point when the MS rates start to increase –rebound onset– and when they stabilize –rebound offset– by doing the following: First, we focused on the MS rate timecourse between 1500-2700 ms after the precue onset, for each observer and each experimental condition. Then we computed the first derivative of the MS rates within this period, and identified where it crossed zero. These specific points were considered candidate rebound onset and offsets. Three independent raters (including one author A.D. and two research assistants) independently identified the inflection points as the start and end points of the MS rate rebounds for each observer and experimental condition. The raters were blind to these conditions, which were presented in a random order.

### Microsaccade timing around the stimulus onset

Precise timings of the MS were analyzed to investigate the effects of temporal attention and expectation on the MS that happen around the stimulus presentation timing. We analyzed the timing of the last MS that occurred before the stimuli presentation (last pre-T1 MS), and the first MS that occurred after the stimuli offset (First post-T1 MS) (Bonneh, Adini, Polat, 2015; Denison et al., 2019). Trials without any MS were excluded from this analysis. We followed the MS timing analysis in the Denison, Yuval-Greenberg and Carrasco (2019) study. In short, for each observer, we first converted all of the MS onsets to z-scores and estimated the kernel density for each precue and expectation condition. Density was evaluated using 300 equally spaced points between −5 and 5, and the median of each kernel was computed to perform further statistical analyses.

### Statistical analysis and visualization

Repeated-measures ANOVA and paired t-tests were performed with R (version 4.2.3; R Core Team, 2023; ezANOVA package Lawrence, 2016), and the nonparametric cluster-based permutation tests were performed using MATLAB. Categorical data were analyzed and visualized using R, whereas time series and kernel densities were investigated using MATLAB. We implemented Bonferroni correction for the cluster-based permutation tests, and Holm-correction for pairwise t-tests to reduce the likelihood of a Type-I error.

## RESULTS

### Microsaccade Rate Dynamics

#### Expectation: Hazard Rate vs Temporal Precision

For hazard rate, we analyzed MS rates from the two lowest temporal precision conditions (uniform and wide) as well as the timing of the rate rebound. For temporal precision, we only analyzed the trials where the targets appeared at the Expected moment. These trials had the same target onsets, but were embedded in sessions with different levels of temporal precision (as reported in Duyar et al., 2024).

We first compared MS rates in trials in which the stimuli appeared Early, Expected, or Late, regardless of attention (**Fig 2A**). This analysis enabled us to compare prolonged prestimulus inhibition vs stimulus-induced inhibition. The cluster-based permutation tests revealed three significant clusters in the inhibitory period, and two during the rebound. In the inhibitory period, observers made less frequent MS in the Early than the Expected trials (1189-1503 ms, p=0.028), and in the Early than Late trials (1239-1639 ms, p=0.007), and in the Expected than Late trials (1415-1663 ms, p=0.039). Subsequently, there were modulations in the rebound, in which MS rates increase back to baseline level. Observers made less frequent MS when the stimuli appeared Late than Early (1681-2017 ms, p=0.003) and than Expected (1739-2037 ms, p=0.007). Overall, strength and timing of inhibition and rebound followed hazard rate.

**Figure 2.**
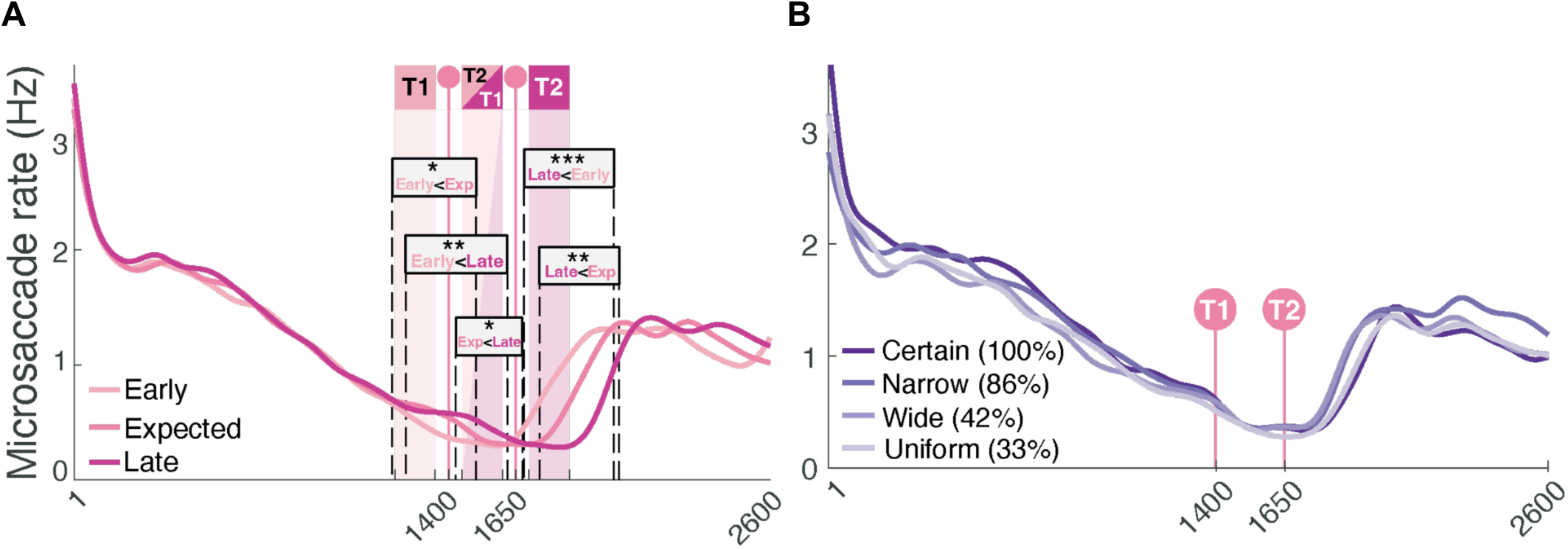
MS rates compared among temporal expectation conditions, regardless of attention locked to precue onset. **A.** MS rate varied with hazard rate. There are three significant clusters in the inhibitory period, where MS rates were higher for delayed stimuli, and two clusters during the rebound, where MS rates were lower with delayed stimuli. Shaded regions indicate Early and Late onsets, and the vertical lines with circles indicate the Expected moment. **B.** MS rates were similar among different levels of temporal precision.*p<0.05; **p<0.01;*p< 0.05; ***p<0.001.

We then compared MS rates among temporal precision levels (**Fig 2B**). Permutation cluster tests revealed no significant differences among the temporal precision levels. MS rate dynamics were similar regardless of temporal precision; they decreased before the stimulus onset, and rebounded after stimulus offset.

We split the data based on attentional precue. For hazard rate (**Fig 3A**), in the Neutral trials (top panel), in the inhibitory period, observers made less frequent MS in the Early than Late (1069-1611 ms, p=0.001) and Expected (1191-1480 ms, p=0.023), and in the Expected than Late (1408-1632 ms, p=0.015) trials. This inhibition was followed by rebound modulations, in which observers made less frequent MS in the Late than Early (1675-2005 ms, p=0.016) and Expected (1729-2011 ms, p=0.002) trials. When T1 was precued (middle panel), in the inhibitory period, observers made less frequent MS in the Early than Late (1249-1568 ms, p=0.020) trials, followed by three clusters during the rebound, and an extra cluster in the post-rebound. During rebound, MS rates were lower in the Expected (1619-1862 ms, p=0.011) and Late (1649-1985 ms, p<0.0001) than Early trials, as well as in the Late than Expected (between 1727-2021 ms, p<0.0001) trials. And in the post-rebound, MS rates were lower in Expected than Late trials (2338-2600 ms, p=0.012). When T2 was precued (bottom panel), the MS rates were similar during the inhibitory period, whereas there were clusters after stimulus offset. During rebound, MS rates were lower in the Late than Early (1688-2009 ms, p<0.0001) and Expected (1747-2035 ms, p=0.001) trials. Subsequently, in the post-rebound, MS rates were lower in the Early than Expected (2233-2492 ms, p=0.002), Early than Late (2313-2542 ms, p=0.021), and in then Expected than Late (between 2390-2600 ms, p=0.016) trials.

**Figure 3.**
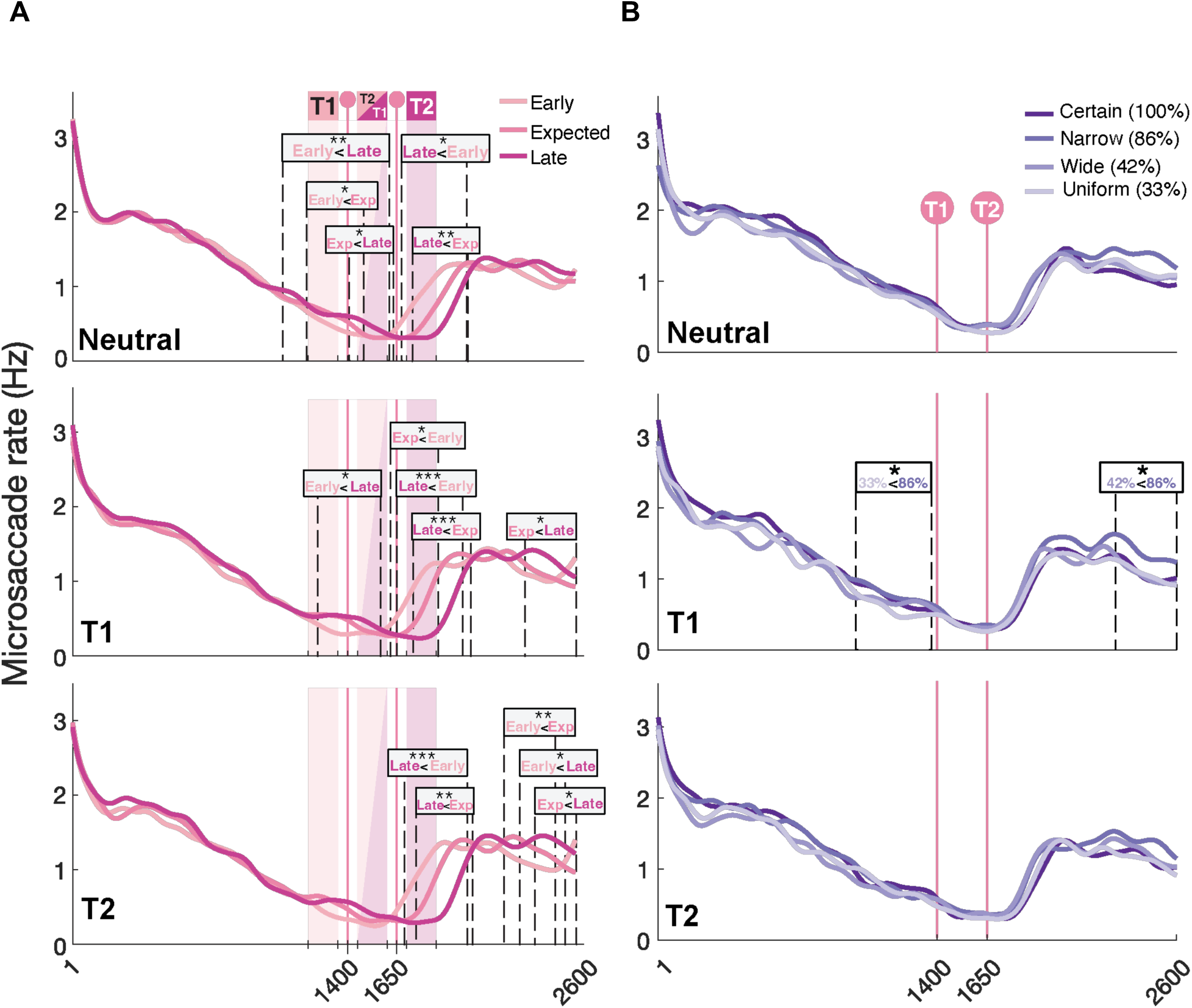
MS rates compared among temporal expectation conditions, separately for each attentional precue (Neutral: top panel, T1-precued: middle panel, T2-precued: bottom panel) trials). **A.** Darker colors indicate a higher hazard rate within the trial. **B.** Darker colors indicate a higher temporal precision.

For temporal precision (**Fig 3B**), there were no significant differences among temporal precision levels, in the Neutral (top panel) or T2 (bottom panel) trials. When T1 was precued (middle panel), two clusters emerged. There were less MS (993-1371 ms, p=0.033) during the inhibitory period in the Uniform (33%) than Narrow (86%) trials. And after the rebound (2293-2600 ms, p=0.045) there were less MS in the Wide (42%) than Narrow (86%) condition.

#### Temporal Attention: Hazard Rate vs Temporal Precision

We compared the attentional conditions by collapsing across hazard rate levels within Wide and Uniform conditions (**Fig 4A**). Cluster-based permutation tests revealed significant differences during the inhibitory period. MS rates were lower for T1 (251-1508 ms, p<0.001) or T2 (599-1303 ms, p=0.003) than Neutral trials. There were similar attentional modulations in the inhibitory period within each hazard rate condition (**Fig 4B**). For Early targets (top panel), MS rates were lower when observers attended to T1 than to Neutral (627-1453 ms, p=0.002), which coincides with the T1 onset. For Expected targets (middle panel), MS rates were lower for T2 (604-1036 ms, p=0.010) and T1 (621-1336 ms, p=0.001) than Neutral. For Late targets (bottom panel), MS rates were lower in T2 (970-1264 ms, p=0.021) and in T1 (987-1306 ms, p=0.015) than Neutral trials. When observers attended to the first target, the modulations coincided with Early T1 onsets (between 1200-1350), but never coincided with any T2 onsets when observers attended to the second target.

**Figure 4.**
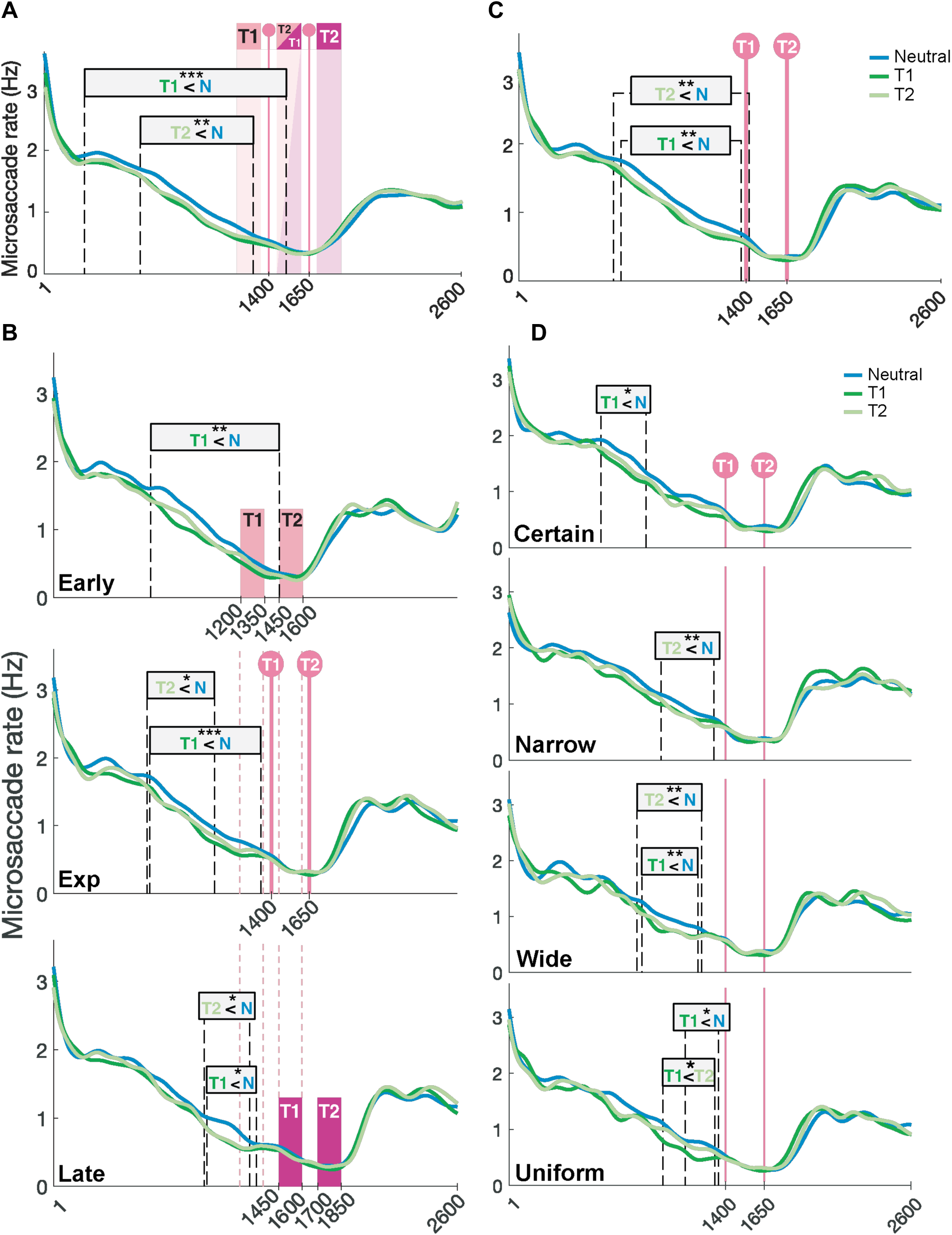
MS rates compared among attentional precue. Blue indicates neutral, green indicates valid attentional precue (dark green: T1-precued, light green: T2-precued). **A-B** includes trials from Wide and Uniform sessions, and **C-D** includes trials in which the stimuli occurred at the expected moment. **A.** MS rates collapsed across Hazard levels. **B.** Attentional precue effects separated by Hazard Rate (Early: top panel, Expected: middle panel, Late: bottom panel). **C.** MS rates collapsed across Precision levels. **D.** Attentional precue effects separated by Precision (Certain: top panel, Narrow: 2^nd^ panel, Wide: 3^rd^ panel, Uniform: bottom panel).

We then compared the attentional conditions by collapsing across temporal precision levels at the expected moment (**Fig 4C**). There were two practically overlapping clusters identified during the inhibitory period. Observers made less frequent MS when attending to T1 (582-1418 ms, p=0.002) or T2 (630-1370 ms, p=0.006) than Neutral. Splitting these data according to temporal precision revealed that similar attentional modulations were present during the inhibitory period within each temporal precision condition (**Fig 4D**). A cluster was identified in the Certain condition (595-886 ms, p=0.022) in which observers made less frequent MS for T1 than Neutral trials. In the Narrow condition, a similar difference emerged between T2 and neutral precue conditions (984-1324 ms, p=0.009). Two clusters practically overlapped in the Wide condition, with lower MS rate in T1 (860-1221 ms, p=0.003) and T2 (828-1247 ms, p=0.006) than in Neutral trials. In the Uniform condition, a similar cluster emerged when attending to T1 than to T2 (994-1330 ms, p=0.010).

We also analyzed the timing and the levels of MS rate rebound. The (three) inter-rater consistency for rebound onset was 91% and for offset was 90% of the trials. In the remaining cases, we included consistently chosen points by two out of three raters. We conducted two-way ANOVAs (precue x hazard rate) on the following parameters:

Rebound onset was affected by hazard (**Fig 5A**, F(2,30)=35.387, p<0.0001, *η^2^*G=0.239), but not by precue (**Fig 5B**, F(2,30)=2.290, p=0.119) or an interaction between these factors (F(4,60)=1.475, p=0.221). Pairwise comparisons revealed that the rebound onset was delayed with stimulus onset (all ps<0.001; Early-Expected: d=0.792, Early-Late: d=1.288, Expected-Late: d=0.659).

**Figure 5.**
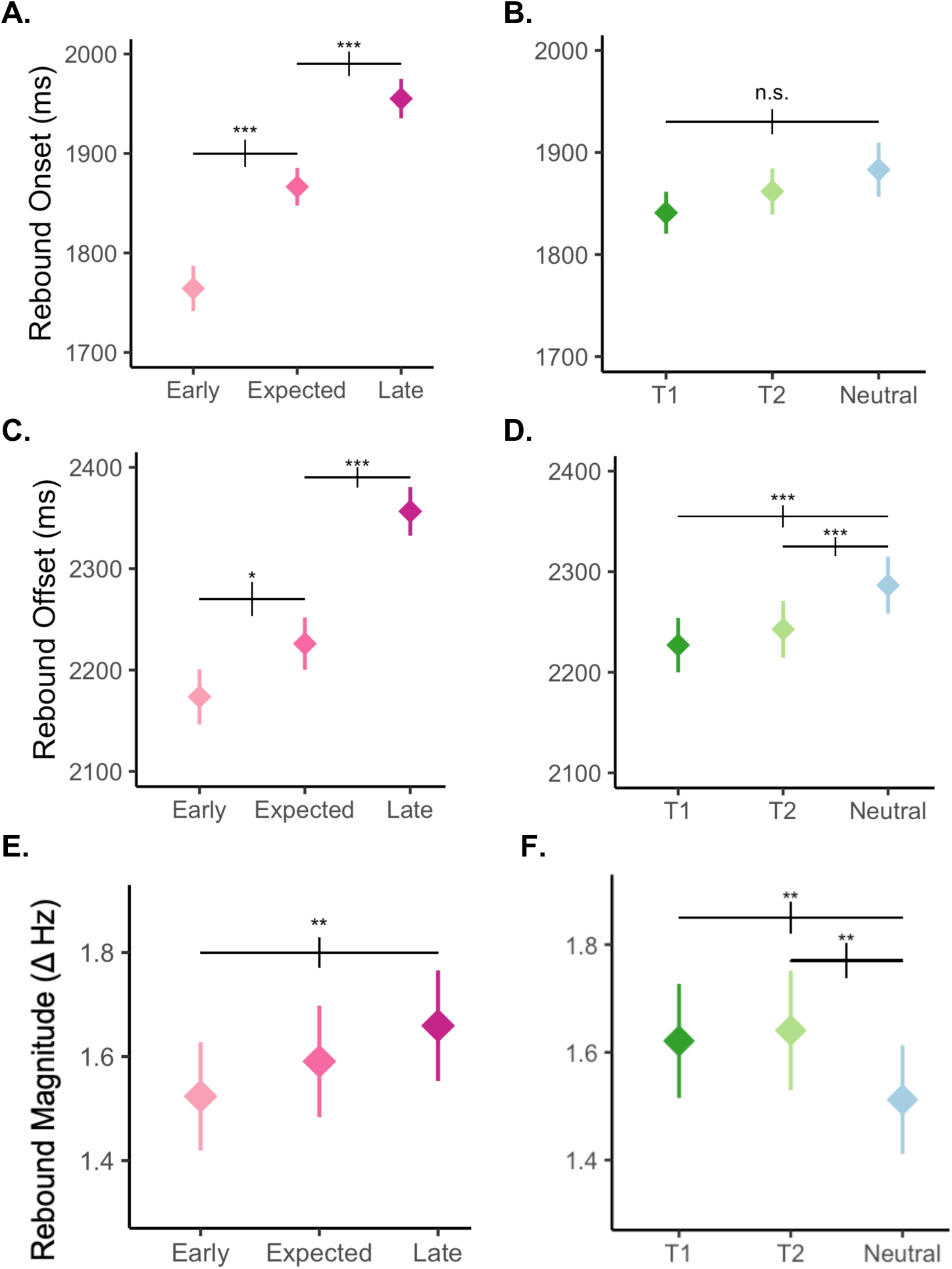
MS rate rebound parameters for Hazard Rate and attentional precue. Left column shows data collapsed across attention, and the right column shows data collapsed across the hazard rate. Error bars represent SEM. **A.** Main effect of Hazard on MS rebound onset. **B.** No significant effect of the attentional precue on MS rebound onset. **C.** Main effect of Hazard on MS rebound offset. **D.** Main effect attentional precue on MS rebound offset. **E.** Main effect of Hazard on MS rebound magnitude. **F.** Main effect attentional precue on MS rebound magnitude.

Rebound offset was affected by hazard (**Fig 5C**, F(2,30)=19.731, p=0.0001, *η*_G_*^2^*=0.162) and precue (**Fig 5D** F(2,30)=11.020, p=0.0003, *η*_G_*^2^*=0.020). Rebound stopped sooner when the stimulus was Early than Expected, (p=0.015, d=0.284) and when it was Expected than Late (p<0.001, d=0.754). Moreover, the rebound stopped sooner for T1 (p<0.001, d=0.309) or T2 (p=0.001, d=0.224) than in the Neutral trials.

Rebound magnitude was affected by hazard (**Fig 5E**, F(2,30)=9.095, p=0.0008, *η*_G_*^2^*=0.006) and precue (**Fig 5F**, F(2,30)=3.969, p=0.0295, *η*_G_*^2^*=0.006). The rebound was smaller for Early than Late targets (p=0.0023, d=0.186). Moreover, the rebound was larger when observers attended to T1 (p=0.019, d=0.152) or T2 (p=0.018, d=0.175) than in the Neutral trials.

Rebound slope was affected by hazard (F(2, 30)=4.036, p=0.028, *η*_G_*^2^*=0.018) and precue (F(2,30)=7.443, p=0.002, *η*_G_*^2^*=0.012). The slope was steeper at Expected than Early (p=0.005, d=0.305) targets, and for T1 (p=0.028, d=0.189) or T2 (p=0.009, d=0.249) than in Neutral trials. Rebound offset MS rate was affected by hazard rate (F(2,30)=4.351, p=0.018, *η*_G_*^2^*=0.002). MS rate was lower at the rebound offset for Early than Late targets (p=0.043, d=0.127). Lastly, there was no main effect or interaction of precue and hazard rate on MS rate for rebound onset (all Fs<1, ps>0.05) or duration (hazard: F(2,30)=2.112, all Fs<1, all ps>0.1).

In sum, with increasing hazard: (i) MS rate inhibition was delayed when collapsing across attentional conditions. This effect was more pronounced in the Neutral than in Attended trials, with the effect being present in T1 but not in T2; (ii) MS rate rebound was delayed and its magnitude increased. But regardless of hazard rate, attention: (i) facilitated MS inhibition; (ii) increased the rebound magnitude, steepened its slope, and hastened its offset.

### Microsaccade Latency Dynamics

To investigate whether the precise timing of the MS are affected by the interaction between temporal attention and expectation, we analyzed the timing of the MS that occurred around the stimulus onset. We analyzed the timing of the last MS that occurred before the stimuli (last pre-T1 MS), and the timing of the first MS that occurred after the stimuli (first post-T1 MS) (Bonneh, Adini, & Polat, 2015; Denison, Yuval-Greenberg & Carrasco, 2019). Analyzing first post-T2 MS yielded the same results because very few MS occurred between T1 and T2.

#### Hazard Rate

Given that target timings varied among trials, we analyzed last pre-T1 and first post-T1 MS latency both relative to precue, as well as to the target onset. We performed two-way (3 hazard rate x 3 precue) ANOVAs on the median timings of the estimated kernel densities.

Relative to precue onset, ANOVA on the last pre-T1 MS latency revealed main effects of the hazard rate (F(2,30)=26.012, p<0.0001, *η*_G_*^2^*=0.112) and attention (F(2,30)=11.875, p=0.002, *η*_G_*^2^*=0.104), but no interaction (F(4,60)=1.284, p=0.293). Pairwise t-tests showed that the last pre-T1 MS was sooner for Early than Expected (p<0.0001, d=0.478), which in turn was sooner than Late (p<0.001, d=0.339) targets (**Fig 6A**), and when observers attended to T1 than to T2 (p=0.017, d=0.216), which in turn was sooner than Neutral (p=0.0001, d=0.542) (**Fig 6B**).

**Figure 6.**
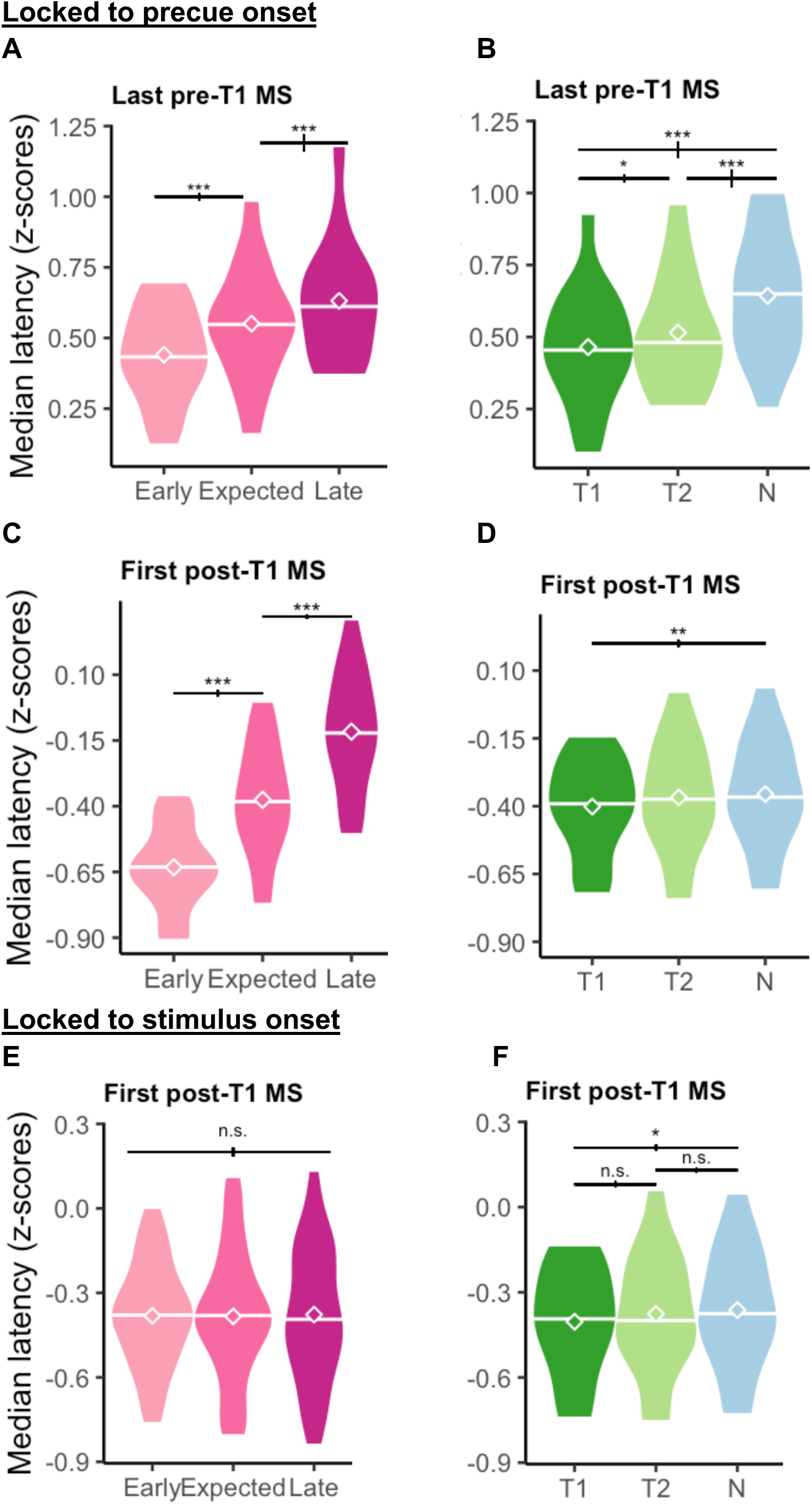
Precise timing of MS that occurred around the stimulus. Diamond icons and the horizontal bars overlaid on the violin plots represent mean and median values, respectively. **A.** Main effect of Hazard on the last pre-T1 MS timing. **B.** Main effect of attentional precue on the last pre-T1 MS timing. **A-D** includes trials locked to precue onset; **E-F** includes trials locked to stimulus onset**. C.** Main effect of Hazard. **D.** Main effect of attentional precue. **E.** Main effect of Hazard. **F.** Main effect of attentional precue.

ANOVA on the first post-T1 MS rebound locked to the precue yielded main effects of hazard rate (F(2,30)=202.788, p<0.0001, *η*_G_*^2^*=0.513) and attention (F(2,30)=4.212, p=0.024, *η*_G_*^2^*=0.009). Pairwise comparisons revealed that the first post-T1 MS were delayed with hazard (**Fig 6C**; all p<0.0001; Early-Expected: d=1.303, Early-Late: d=2.596, Expected-Late: d=1.131). Moreover, the first post-T1 MS were earlier when T1 was precued than in Neutral trials (p=0.008, d=0.158) (**Fig 6D**).

Relative to target onset, ANOVA on last pre-T1 MS latency revealed main effects of hazard rate (F(2,30)=144.793, p<0.0001, *η*_G_*^2^*=0.449) and attention (F(2,30)=12.026, p=0.0001, *η*_G_*^2^*=0.103), but no interaction (F(4,60)=1.508, p=0.231). Pairwise t-tests showed that the time window between the last pre-T1 MS and the T1 onset was the shortest for Early than Expected (p<0.0001, d=1.022), which in turn was shorter than for Late (p<0.0001, d=1.052). Similarly, it was shorter when attending to T1 than T2 (p=0.050, d=0.147), which in turn was shorter than for Neutral (p<0.001, d=0.423) trials. These findings are in the same direction as the MS timing locked to the precue.

A 2-way ANOVA on the median first post-T1 MS timing showed a main effect of precue (F(2,30)=0.034, p=0.034, *η*_G_*^2^*=0.006), but not of hazard (F(2,30)<1, p>0.1; **Fig 6E**). Post-T1 MS was sooner when T1 was precued than in Neutral trials (p=0.018, d=0.201; **Fig 6F**).

### Temporal Precision

We performed 2-way (4 precision x 3 precue) ANOVAs on the median timings of the last pre-T1 and first post-T1 MS. On both last pre-T1 and first post-T1 MS timings, there was a main effect of attention (last pre-T1 **Fig 7B**: F(2,26)=10.111, p<0.001, *η*_G_*^2^*=0.067; first post-T1 **Fig 7D**: F(2,26)=7.847, p=0.002, *η*_G_*^2^*=0.017), but not of precision (last pre-T1: F<1, p>0.1; first post-T1: F<1, p>0.1) or their interaction (last pre-T1 **Fig 7A**: F(6,78)=1.271, p>0.1; first post-T1 **Fig 7C**: F(6,78)=1.563, p>0.1). Post-hoc pairwise t-tests showed that the last pre-T1 MS was earlier in T1 (p<0.001, d=0.599) and T2 (p<0.001, d=0.506) than in Neutral trials. First post-T1 MS were earlier when attending to T1 than T2 (p=0.003, d=0.273) or Neutral (p=0.006, d=0.276) trials.

**Figure 7.**
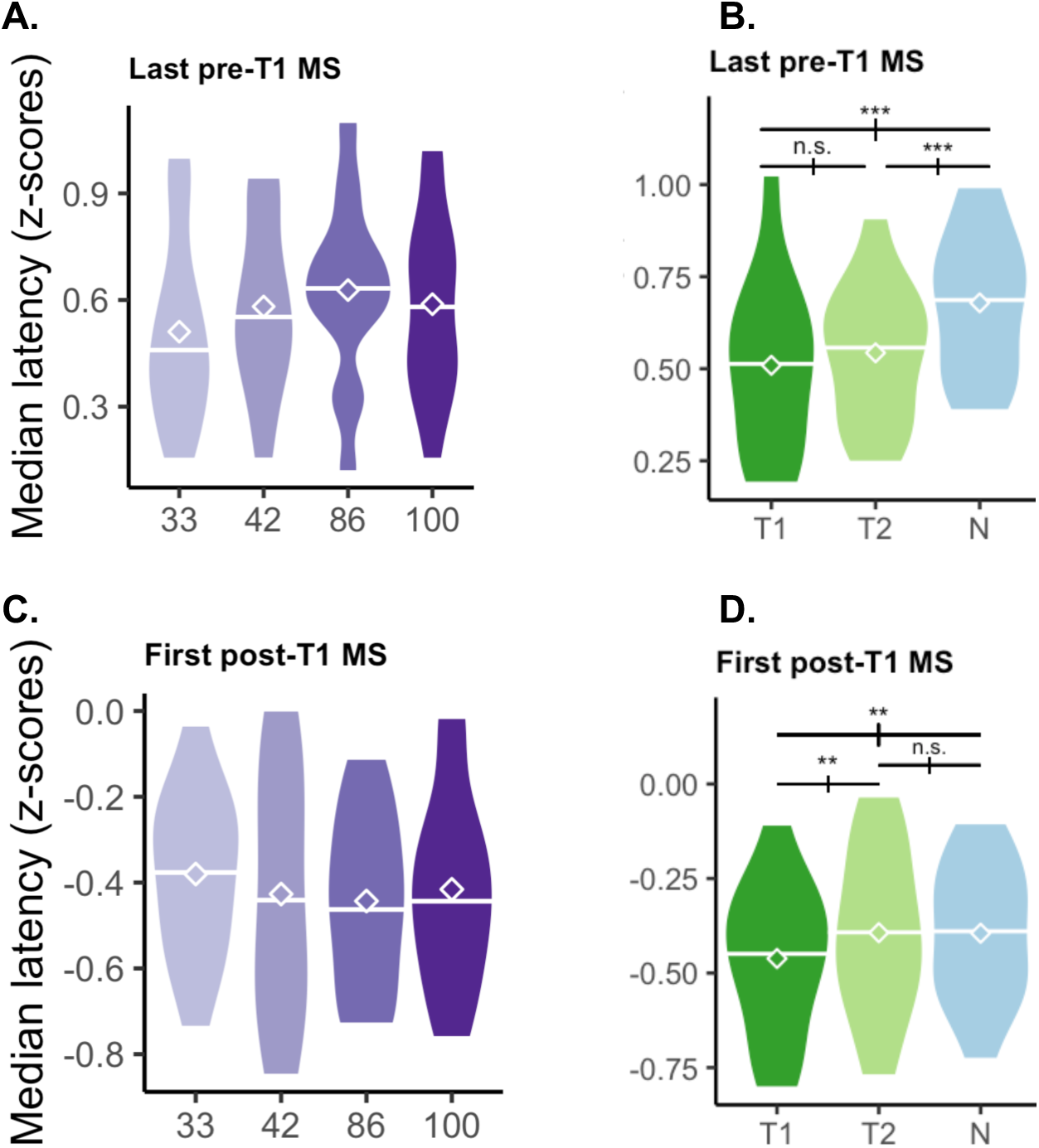
Precise timing of MS that occurred around the stimulus. **A.** No effect of Temporal Precision on the last pre-T1 MS timing. **B.** Main effect of attentional precue on the last pre-T1 MS timing. **C.** No effect of Temporal Precision on the first post-T1 MS timing. **D.** Main effect of attentional precue on the first post-T1 MS timing.

## DISCUSSION

Different temporal structures shape temporal expectations, influencing perception, action and attention through overlapping yet distinct mechanisms, such as modulations in neural signal strength, latency and synchrony (review: Nobre & van Ede, 2017). The brain forms temporal expectations through distinct neural correlates, which are integrated into behavioral responses, based on its temporal structure (Coull et al., 2000; Tal-Perry & Yuval-Greenberg, 2020; Grabenhorst et al., 2021). Hazard rate–mounting expectations through time passage–and temporal precision–based on probabilistic distributions–differentially influence auditory neural responses (Todorovic & Auksztulewicz, 2021) and interact with temporal attention distinctly when modulating visual performance (Duyar et al., 2024). Here, explored potential dissociations between perception and action and examined whether they can be distinguished at the oculomotor level.

We focused on the temporal dynamics of the push-pull mechanisms of MS rate signature across three stages: (1) pre-stimulus anticipatory reduction, (2) post-stimulus reflexive exogenous inhibition, and (3) rebound, which enabled us to disentangle endogenous components driven by anticipatory processes from the exogenous components triggered by external stimuli.

### MS rate signature is aligned with the hazard rate, but not with precision

We analyzed MS rate signatures across varying hazard rates and across different precision levels. As the stimuli were delayed and the hazard rate increased, prestimulus anticipatory reduction was prolonged, and post-stimulus reflexive exogenous inhibition and the corresponding rebound were delayed. And notably, the post-stimulus inhibition was stronger than the anticipatory reduction, resulting in lower MS rates in Early than Late trials (**Fig 2A**). In contrast, MS rate signatures were similar across temporal precision levels (**Fig 2B**).

### Temporal attention modulates hazard rate and temporal precision in an opposite manner during MS rate decrease

We analyzed MS rate signature across varying hazard rates and across different precision levels as a function of attention. In the neutral condition, MS rates were lower when the target was early during the reduction/inhibition, and the corresponding rebound occurred earlier; however, no such effect was observed for temporal precision (**Fig 3A and B, top panels**). When observers attended to T1, prestimulus reduction was stronger, leading to an attenuated reflexive inhibition and resulting in only one cluster during the inhibitory period, highlighting the difference between Early and Late trials. As a result, the distinctions between the hazard rate levels that were present in Neutral were diminished when attending to T1 and disappeared when attending to T2 (**Fig 3A, middle and bottom panels**). In contrast, precision levels were distinct during the pre-stimulus period when observers attended to T1 (**Fig 3B, middle panel**). Specifically, microsaccades occurred less frequently in the pre-stimulus period under lower (33%) than higher (86%) temporal precision. Because hazard rate depends on stimulus onset, attentional interaction’s emergence in the post-stimulus period is expected.

However, the prestimulus emergence of the precision effect is notable, especially given its late effect on auditory-evoked MEG responses compared to hazard rate (Todorovic & Auksztulewicz, 2019). Furthermore, our findings indicate that temporal precision information is integrated into eye movement planning primarily during attentional deployment in the anticipatory period.

### Temporal attention modulates hazard rate and temporal precision similarly after rebound

Notably, we found that post-rebound differences emerged with attention, both for hazard rate (**Fig 3A, middle and bottom panels**) and for temporal precision (**Fig 3B, middle panel**). These differences were in the same direction, such that a higher probability of the stimulus onset was associated with a higher MS rate in the post-rebound. This similarity suggests a potential common mechanism for integrating the attended probabilistic temporal information irrespective of the temporal structure from which the probability is derived. Such post-stimulus processing commonality among hazard rate and temporal precision was observed in gain modulation in primary auditory cortex and superior temporal gyrus (Todorovic & Auksztulewicz, 2019).

### Perception-action parallelism is observed for the hazard rate, but not for temporal precision

Temporal attention facilitates pre-stimulus anticipatory MS reduction when stimulus timing is predictable (Denison et al., 2019; Palmieri et al., 2023), as well as under low temporal precision (Duyar & Carrasco, 2024). Here we found that attentional modulation can extend beyond T1(**Fig 4A**). Remarkably, this modulation coincides with early stimuli onsets, for which hazard is the lowest (**Fig 4B**), within the window with largest attentional benefits on performance (Duyar et al., 2024). Together, these results suggest a parallelism between performance and oculomotor behavior. We did not observe such a parallelism for temporal precision. Attentional modulation occurred earlier for Certain than all uncertain trials (**Fig 4D**).

### Temporal attention and hazard rate operate independently during the post-stimulus period

We quantified both rebound temporal shift and magnitude in hazard rate. The MS rate rebound is not merely a reflexive release from prior suppression; it is delayed following an oddball event (Valsecchi, Betta & Turatto, 2007; Widmann, Engbert & Schroger, 2014) and with task difficulty due to prolonged sensory evidence accumulation (Ezzo et al., 2025). Its duration also increases as a function of reaction times (Betta & Turatto, 2006), while its magnitude diminishes following an oddball event (Valsecchi & Turatto, 2009). Moreover, the magnitude of the rebound is linked to activity in the frontal eye fields (Hsu, Chen, Tseng & Wang, 2021; Fernández, Hanning, Carrasco, 2023). Unlike the interaction between temporal attention and hazard rate on performance, we observed no evidence for an interaction between them on MS rate rebound. In particular, rebound onset was delayed with hazard (**Fig 5A**), but not with attention (**Fig 5B**). Rebound offset was also delayed with hazard (**Fig 5C**), but attention hastened it (**Fig 5D**), so that post-stimulus MS rate reached baseline level faster with attention. We also found that rebound magnitude increased with both hazard (**Fig 5E**) and attention (**Fig 5F**). Overall, these findings suggest that temporal attention and expectation operate independently during the post-stimulus period. The delay in rebound onset and offset with hazard rate primarily indicates a temporal shift locked to stimulus timing, whereas attention hastened rebound offset, expediting return to baseline rate without affecting its onset. Furthermore, the increase in rebound magnitude with both hazard and attention may reflect their separate contributions to ongoing cognitive control and decision making during the post-stimulus window.

Precise timing of MS around stimulus onset allows for quantification of oculomotor freezing and the subsequent release, revealing fine-grained temporal dynamics of perceptual and cognitive processes (Bonneh et al., 2015; Yablonski, Polat, Bonneh & Ben-Shachar, 2016). When the stimulus timing is certain, the last MS before stimulus onset and the first MS after stimulus onset occur earlier with temporal attention (Denison et al., 2019).

### Perception-action dissociation for the hazard rate

Effects of temporal attention and hazard rate interact on performance (Duyar et al., 2024), but we found not such an interaction on the precise timing of MS. The oculomotor freezing onset—indexed by indexed by last pre-T1 MS— was delayed with the hazard rate, as the stimulus was delayed (**Fig 6A**). But regardless of hazard rate, attention speeded oculomotor freezing (**Fig 6B**). Correspondingly, release from freezing, indexed by the first post-T1 MS was delayed with hazard rate (**Fig 6C**) and shifted earlier with attention (**Fig 6D**).

### Upcoming MS are scheduled in advance

To differentiate the effects of MS timing from those of stimulus latency, we analyzed MS timing relative to the stimulus within each trial. This analysis revealed that the release from inhibition was consistent regardless of hazard rate, with the first post-stimulus MS occurring after a similar duration following stimulus onset (**Fig 6E**). In comparison, attentional effects on the release from inhibition were still present relative to stimulus onset, such that the release from inhibition occurred earlier in the attended trials as compared to Neutral (**Fig 6F**). This provides evidence for the preparatory scheduling of MS in advance (Badde et al., 2020). Hazard rate is stimulus-dependent and dynamically updated as the stimulus is delayed. Consequently, the release from inhibition cannot be pre-planned with respect to hazard rate. The release timing was unaffected by hazard rate, which is inherently tied to immediate, transient changes. In contrast, attentional allocation occurs earlier within the trial, enabling planning of the inhibition release and advance scheduling of MS. Overall, we conclude that release from inhibition can be governed by endogenous anticipatory rather than transient processes.

### Perception-action dissociation for temporal precision

Although temporal attention benefits on visual performance increase with temporal precision (Duyar et al., 2024), we found precise timing of MS is independently and consistently advanced with attention. Regardless of temporal precision, the oculomotor freezing onset—the last pre-T1 and first post-T1 MS were consistent (**Fig 7A & 7C**), but started (**Fig 7B**) and released (**Fig 7D**) earlier with attention.

The temporal scale of temporal precision and hazard rate differs. Understanding how two probabilistic structures are represented may provide valuable insights regarding atypical conditions, such as schizophrenia, attention deficit hyperactivity disorder, autism, and Parkinson (e.g., Aitkin, Santos & Kowler, 2013; González-Gadea et al., 2015; Dankner et al., 2017; Martin et al., 2017; Cannon et al., 2021; Coull & Giersch, 2022; Degos, Pouget & Missal, 2022) in which temporal expectations are altered. Revealing similarities and distinctions between different types of temporal expectations is important for improving cognitive models, which can be useful in developing diagnostic and interventional tools. In this context, MS serves as a valuable noninvasive biomarker.

In conclusion, expectations based on hazard rate are represented within the oculomotor system, with probabilistic information through time passage contributing to fixational stability. In contrast, we found no evidence that temporal expectation based on precision is represented in the oculomotor system, except when temporal attention is allocated. We observed action-perception dissociations for both types of expectation on the precise timing of MS, alongside parallel instances in which attentional modulations on MS coincided with performance benefits with increasing hazard rate. Our study provides insights into the endogenous and stimulus-driven components of oculomotor dynamics, demonstrating how temporal attention and distinct types of expectation work together to enhance fixational stability.

## Acknowledgments

We thank Lucy Gardiner and Sherry Ye for their contributions, as well as Rania Ezzo, Mariel Roberts and other members of the Carrasco Lab for providing helpful comments on the manuscript. This research was funded by the National Eye Institute (R01-EY019693 to M.C.).

The authors declare no competing interest.

## Notes

### Competing Interest Statement

The authors have declared no competing interest.

